# NMA1982 is a Novel Phosphatase and Potential Virulence Factor in *Neisseria meningitidis*

**DOI:** 10.1101/2023.05.23.541968

**Authors:** Shuangding Wu, Mathieu Coureuil, Xavier Nassif, Lutz Tautz

## Abstract

Protein phosphorylation is an integral part of many cellular processes, not only in eukaryotes but also in bacteria. The discovery of both prokaryotic protein kinases and phosphatases has created interest in generating antibacterial therapeutics that target these enzymes. NMA1982 is a putative phosphatase from *Neisseria meningitidis*, the causative agent of meningitis and meningococcal septicemia. The overall fold of NMA1982 closely resembles that of protein tyrosine phosphatases (PTPs). However, the hallmark C(X)_5_R PTP signature motif, containing the catalytic cysteine and invariant arginine, is shorter by one amino acid in NMA1982. This has cast doubt about the catalytic mechanism of NMA1982 and its assignment to the PTP superfamily. Here, we demonstrate that NMA1982 indeed employs a catalytic mechanism that is specific to PTPs. Mutagenesis experiments, transition state inhibition, pH-dependence activity, and oxidative inactivation experiments all support that NMA1982 is a genuine phosphatase. Importantly, we show that NMA1982 is secreted by *N. meningitidis*, suggesting that this protein is a potential virulence factor. Future studies will need to address whether NMA1982 is indeed essential for *N. meningitidis* survival and virulence. Based on its unique active site conformation, NMA1982 may become a suitable target for developing selective antibacterial drugs.

## Introduction

Protein phosphorylation is considered a universal mechanism to regulate protein function during most cellular processes [1, 2]. About one third of all eukaryotic proteins are phosphorylated, predominantly on serine, threonine, and tyrosine [3]. In bacteria, phosphorylation on aspartate and histidine is well characterized as part of the two-component system [4]. However, bacterial proteins are also phosphorylated on serine, threonine, and tyrosine [5, 6]. Interestingly, phosphorylation on tyrosine is much more abundant in bacterial proteins (9%), relative to serine and threonine, than it is in humans (1.8%) [7]. Both protein tyrosine kinases (PTKs) and protein tyrosine phosphatases (PTPs) have been isolated from bacteria [8-10] and have been proposed as anti-bacterial drug targets [10-12].

The PTP superfamily contains various subfamilies with the two largest being the classical PTPs, which only dephosphorylate phosphotyrosine (pTyr), and the dual-specificity phosphatases (DUSPs), which in addition to pTyr can also dephosphorylate serine, threonine, or non-protein substrates such as phospholipids or RNA [13, 14]. All PTPs have a cysteine-based catalytic mechanism (**Fig 1**), which involves the nucleophilic attack on the phosphate group by the catalytic cysteine thiolate, resulting in the cleavage of the phosphoester bond and the formation of a phospho-enzyme intermediate [15]. This first catalytic step is followed by another nucleophilic attack of a coordinated water molecule, resulting in the hydrolysis of the sulfur-phosphorus bond, the release of inorganic phosphate, and the recovery of the enzyme. The catalytic cysteine as well as the invariant arginine, which stabilizes the transition state, are part of the PTP signature motif C(X)_5_R that forms the phosphate-binding loop, also known as the P-loop. Catalysis is further assisted by an aspartic acid residue that functions both as the catalytic acid, donating a proton to the leaving group in the first step, and the catalytic base, activating the water molecule in the second step of the PTP reaction. This aspartic acid is part of the WPD-loop, named after the conserved tryptophan–proline–aspartate (WPD) motif in all classical PTPs. In DUSPs, only the aspartic acid is conserved; hence, this loop is also referred to as the D-loop.

**Fig 1.**
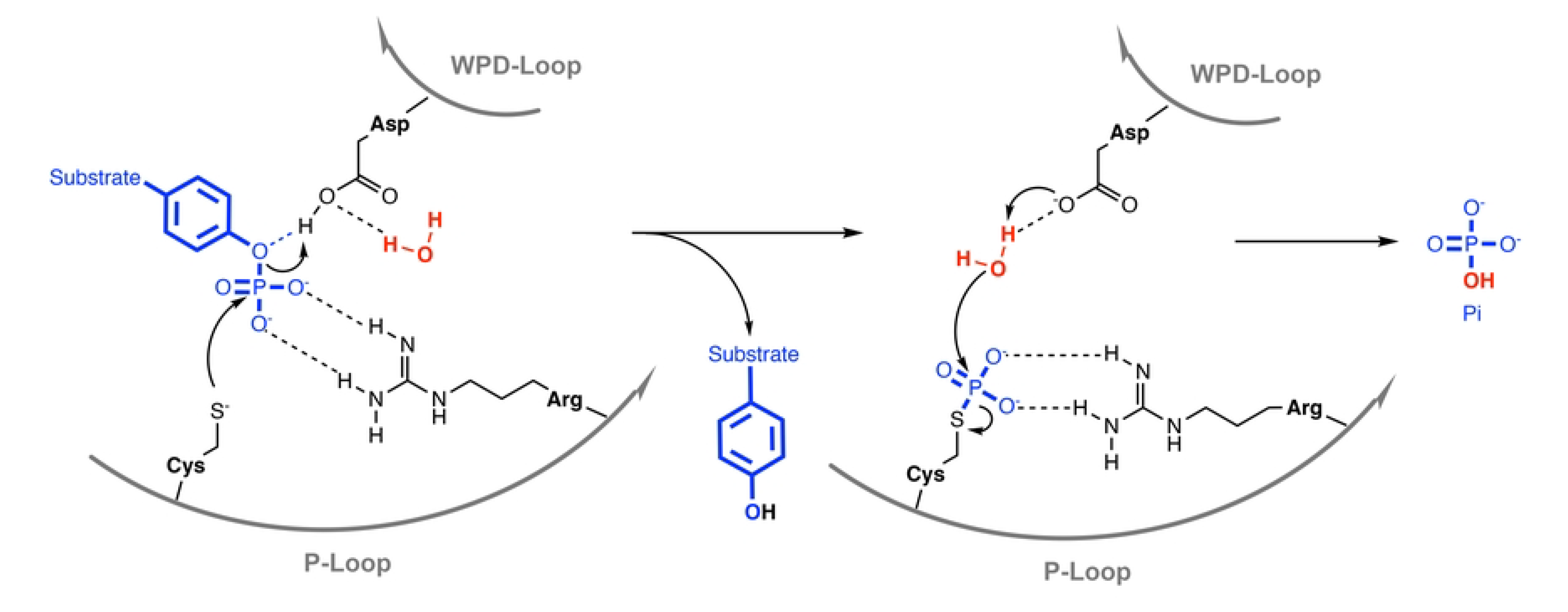
PTP catalytic mechanism. See text for details.

The WPD-loop fluctuates between an “open” conformation, allowing substrate binding, and a “closed” conformation, necessary for catalysis. Decreased mobility of the WPD-loop can result in substantially lower catalytic activity, as shown by mutational studies [16, 17] as well as allosteric inhibitors that decrease WPD-loop flexibility [18].

*Neisseria meningitidis* is a human pathogen responsible for causing meningococcal diseases such as meningitis and meningococcal septicemia [19]. Despite established vaccination strategies, *N. meningitidis* continues to be an important cause of mortality and morbidity in newborns and children in both developed and underdeveloped countries [20]. Unlike viral meningitis, meningococcal meningitis can potentially be fatal within 24 h after the first symptoms appear [21]. While antibiotics typically provide an effective treatment, cases of *N. meningitidis* antibiotic resistance have been reported [22]. Thus, the identification of novel drug targets that can be exploited for the treatment of meningitis is an unmet medical need. Previously, we reported the crystal structure of the *N. meningitidis* protein NMA1982, analysis of which suggested that its overall fold shows strong similarity to the DUSP subfamily of PTPs [23]. However, the P-loop containing the hallmark C(X)_5_R PTP signature motif is shorter by one amino acid in NMA1982, forming a C(X)_4_R motif. This disparity has cast doubts about the catalytic mechanism of NMA1982 and the assignment of NMA1982 to the PTP superfamily. Here, we show that NMA1982 indeed uses a PTP-like catalytic mechanism, demonstrating that the shorter C(X)_4_R motif is suitable to perform phosphatase catalytic function. Moreover, using pH-dependence, small molecule transition state inhibition, and oxidative inactivation experiments, we confirm that NMA1982 behaves like a typical PTP. Finally, we show that NMA1982, like other virulence factors, is secreted by *N. meningitidis*, suggesting that this phosphatase may be important for bacterial infection.

## Materials and Methods

### Site-directed mutagenesis, protein expression and purification

Utilizing our previously described pSpeedET plasmid containing the gene encoding NMA1982 [23], a QuickChange site-directed mutagenesis kit (Stratagene) was used for generating the NMA1982 mutants D71A, C95S, R100A, R136A, and R136A-D71A. Mutations were confirmed by DNA sequencing. The plasmids were transformed into Rosetta 2 (DE3) competent cells. Protein was expressed in LB media by induction with 0.1 mM Isopropyl β-D-1- thiogalactopyranoside (IPTG). Wild-type and mutant NMA1982 proteins were purified as previously described [23]. The purity of the proteins was confirmed to be >95% using SDS-PAGE and Coomassie staining. Differential scanning calorimetry (DSC) was used to confirm that all proteins were folded. Recombinant VHX, VHR, PTP1B, LMPTP-A, and LMPTP-B were expressed and purified as described previously [24, 25]. Recombinant MKP-1 was purchased from Upstate. Recombinant CD45 and SHP1 were purchased from Biomol.

### Michaelis-Menten kinetic PTP assays

NMA1982 phosphatase activity was measured using a standard fluorescence intensity phosphatase assay utilizing 6,8-difluoro-4-methylumbelliferyl phosphate (DiFMUP; Invitrogen) as the substrate [26]. The NMA1982-catalyzed hydrolysis of DiFMUP was assayed at room temperature in a 60 μL 96-well format reaction system in 50 mM Bis-Tris, pH 6.0 assay buffer containing 1.7 mM dithiothreitol (DTT) and 0.005% Tween-20. Various fixed concentrations of DiFMUP (800, 400, 200, 100, 50, 25, 12.5, 6.25, 3.125, 1.5625 µM) were added to recombinant wt NMA1982 (11 µM) or NMA1982 mutants D71A (13 µM), C95S (60 µM), R100A (60 µM), R136A (3.8 µM), or R136A-D71A (4.8 µM). The initial rates were determined from the slopes of the fluorescence emission readings over 30 min using an FLx800 micro plate reader (Bio-Tek Instruments, Inc.) with an excitation wavelength of 360 nm and an emission wavelength of 460 nm. The nonenzymatic hydrolysis of the substrate was corrected by measuring the control without addition of enzyme. The Michaelis-Menten constants (*K*_m_) and turnover numbers (*k*_cat_) were determined by fitting the data to the Michaelis-Menten equation, using non-linear regression and the program Prism (v9, GraphPad Software, Inc.). A similar assay setup was used for Michaelis-Menten kinetic assays with PTP1B, VHX, VHR, MKP-1, CD45, SHP1, LMPTP-A, and LMPTP-B.

### Orthovanadate inhibition of NMA1982

NMA1982 phosphatase activity was measured using the DiFMUP kinetic assay system described above. NMA1982 (15 µM) was preincubated with various fixed concentrations of sodium orthovanadate (Sigma; 1666, 833, 416, 208, 104, 52, 26, 13, 6.5, 3.25, 1.62, 0 µM) for 10 min at room temperature. The reaction was initiated by the addition of the substrate (DiFMUP; 35 µM). The IC_50_ value of orthovanadate for NMA1982 was determined from the initial rates using the program Prism (v9, GraphPad Software, Inc.) as described previously [26].

### NMA1982 oxidation assay

NMA1982 (22 µM) was incubated with various fixed concentrations of H_2_O_2_ (Sigma; 10000, 2000, 400, 80, 16, 3.2, 0 µM) in 300 µL assay buffer (50 mM Bis-Tris pH 6.0, 0.005% Tween-20) on ice for 1 or 2 h. Samples were then either subjected to DTT treatment (10 mM) for 20 min at room temperature or no treatment before DiFMUP (200 µM final concentration) was added to start the reaction (30 min). Fluorescence intensity was measured and initial rates determined as described above.

### NMA1982 activity vs. pH assay

NMA1982 (22 µM) was assayed at room temperature in a 60 μL 96-well format reaction system in 50 mM Bis-Tris buffer (adjusted to pH 5.5, 6.0, 6.5, 7.0, or 7.5) containing 1.7 mM DTT, and 0.005% Tween-20. DiFMUP (200 µM final concentration) was added to start the reaction (30 min). Fluorescence intensity was measured and initial rates determined as described above.

### NMA1982 activity vs. temperature assay

The NMA1982-catalyzed hydrolysis of DiFMUP was assayed at various fixed temperatures (22, 37, 50, and 80 °C) in 50 mM Bis-Tris pH 6.0 assay buffer containing 1.7 mM DTT and 0.005% Tween-20. NMA1982 (22 µM) was incubated with DiFMUP (200 µM final concentration) for 30 min in reactions tubes at the specified temperature, before transferred to a 96-well plate for fluorescence measurements. The fluorescence emission was determined at two time points, 0 and 30 min, using an FLx800 micro plate reader (Bio-Tek Instruments, Inc.) with an excitation wavelength of 360 nm and an emission wavelength of 460 nm. The nonenzymatic hydrolysis of the substrate was corrected by measuring the control without addition of enzyme. NMA1982 activity of each sample was determined from the difference in emission intensity between the 0- and 30-min time points.

### NMA1982 antibodies

Polyclonal rabbit anti NMA1982 antibodies were generated by Abnova (Taiwan). Full-length recombinant NMA1982 protein (6 mg/mL) was used for rabbit immunization. Antibody performance from two batches (Lot #11229 and Lot #11230) was tested against recombinant His-NMA1982 (**Fig S1**). Anti NMA1982 antibodies also recognized the NMA1982 ortholog NMV0640.

### Bacterial strains and mutagenesis

The model organism *N. meningitidis* 8013 was used to engineer a strain lacking the NMA1982 ortholog NMV0640. NEM8013 is a piliated and capsulated Opa-variant of the serogroup C strain 2C4.3. The *NMV0640* gene was disrupted by inserting an aphA-3 cassette (kanamycin resistance) in the coding region using PCR overlap and the following primers: Upstream to NMV0640, 5’-GCCGTCTGAAACTCGTGGAACGTCAAATCC-3’ and 5’- CAGCTCATTACTGTTTGATTTGGGCGAAG-3’; aphA-3 cassette, 5’- CAAACAGTAATGAGCTGATTTAACAAAAATTTAACGCG-3’ and 5’- GACGGGATATCAGAAGAACTCGTCAAGAAGGCG-3’; downstream to NMV0640, 5’- TCTTCTGATATCCCGTCCTTGCCTATTG-3’ and 5’- TTCAGACGGCGCAGACAAACAAGCAGGACA-3’. Insertion of the aphA-3 cassette was verified by PCR and the lack of protein expression was confirmed by SDS-Page and Western blotting.

### Preparation of supernatant and analysis

Wild type 2C4.3 strain and its derivative lacking NMV0640 were grown in 50 mL of Dulbecco’s Modified Eagle Medium (DMEM) for 2 h (from OD_600_ = 0.05 to OD_600_ = 0.2). Bacteria were centrifuged for 15 min at 4,000 rpm and the resulting media were filtered (0.22 µm pore size, Filter System, Corning). Supernatants were concentrated 200x using an UltraCell-3K column (Amicon). The resulting fraction contained proteins and potentially outer membrane vesicles. 10 µg of a whole cell lysate and 5 µL of concentrated supernatant were studied by SDS-Page and immunoblotting using the following antibodies: polyclonal rabbit anti NMA1982/NMV0640 (#11230), monoclonal mouse anti-PilE (clone SM1, generously provided by Dr. Mumtaz Virji), monoclonal mouse anti-RMP4 (Coureuil/Nassif laboratory), monoclonal mouse anti-nicotinamide adenine dinucleotide phosphate (NADP) glutamate dehydrogenase (Coureuil/Nassif laboratory), and secondary antibody anti-IgG (anti-mouse or anti-rabbit, Invitrogen #31430 and #31460) coupled to Horse Radish Peroxidase.

## Results

### 3D Structural comparison of NMA1982 with human PTPs

Using pairwise Dali [27] structural alignment of the crystal structures of NMA1982 and human DUSPs, we found that cyclin-dependent kinase inhibitor 3 (CDKN3), also known as kinase-associated phosphatase (KAP), is the most similar human phosphatase to NMA1982, with an amino acid identity of 15% and a Dali Z-score of 14.9 (**Fig 2a**). Importantly, several conserved loops and amino acid residues critical for phosphatase activity [13] could be identified in NMA1982 (**Fig 2b**): *1)* The phosphate-binding loop (P-loop), which forms the center of the active site, is present in NMA1982, albeit shorter by one amino acid. In NMA1982 the putative catalytic cysteine (C95) and invariant arginine (R100), which both align very well with the corresponding residues in CDKN3, form a C(X)_4_R motif instead of the typical C(X)_5_R PTP signature motif. *2)* The general catalytic acid/base aspartate-containing WPD-loop is also present in NMA1982. D71 putatively serves the role of the catalytic acid/base. Notably, the WPD-loop in the NMA1982 crystal structure adopts an atypically open conformation (with D71 far removed from the active site), which also has been observed for several human PTPs [28]. *3)* NMA1982 features a typical PTP E-loop, which contains a conserved glutamic acid that coordinates the side chain of the invariant arginine in the P-loop and positions it for phosphate binding [13]. Indeed, in the NMA1982 crystal structure, the side chain carboxylic acid of E53 forms a salt bridge with the guanidinium group of R100. Collectively, our comparison with human DUSPs confirms that amino acids and loops critical for phosphatase activity are present in NMA1982.

**Fig 2.**
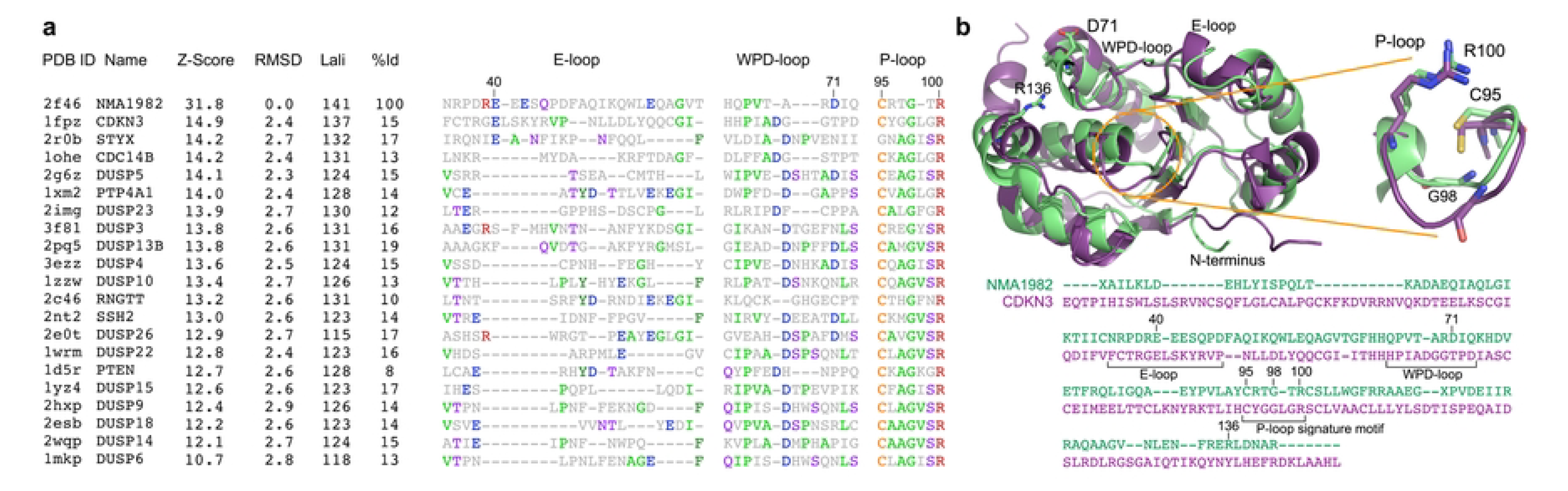
Structural comparison of NMA1982 with human PTPs. **(a)** Pairwise Dali structural alignment of NMA1982 with similar human PTPs. Conserved loops, containing invariant amino acids, are shown. The most frequent amino acid type is colored in each column. Residue numbers correspond to NMA1982. Protein Data Bank (PDB) IDs, Z-Scores as calculated by the DALI server, root mean square deviation (RMSD) between NMA1982 and each homolog structure, length of alignment (Lali), and percent amino acid identity (%Id) are listed. **(b)** Structural alignment of NMA1982 (PDB ID 2f46; green) with its most similar human PTP, CDKN3 (PDB ID 1fpz; purple). Conserved PTP residues in P-loop, WPD-loop, and E-loop are highlighted in stick representation. Residue numbers correspond to NMA1982.

### Assessing NMA1982 phosphatase activity using Michaelis-Menten kinetics

We previously showed that NMA1982 has phosphatase activity using *p*-nitrophenyl phosphate as a pTyr surrogate substrate in a colorimetric assay [23]. To assess the catalytic parameters of NMA1982 more precisely in a kinetic experiment, we adapted a standard, continuous, fluorescence-based phosphatase assay using DiFMUP as the substrate [26, 29]. Using this assay, we performed a Michaelis-Menten experiment to determine the Michaelis-Menten constant and turnover number for NMA1982 (**Fig 3a**). The calculated *K*_m_ value of 35 μM was well in the range of the *K*_m_ values for human PTPs that we assayed under similar conditions, including the DUSPs VHX (*K*_m_ = 11 µM), VHR (*K*_m_ = 39 µM), and MKP-1 (*K*_m_ = 40 µM); the classical PTPs PTP1B (*K*_m_ = 4 µM), CD45 (*K*_m_ = 43 µM), and SHP1 (*K*_m_ = 52 µM); as well as the low-molecular weight PTPs LMPTP-A (*K*_m_ = 63 µM) and LMPTP-B (*K*_m_ = 218 µM). Thus, the Michaelis-Menten kinetic data suggest that the standard PTP substrate DiFMUP is similarly well accepted by NMA1982 and human PTPs. The turnover number for the NMA1982-catalyzed DiFMUP reaction was calculated to *k*_cat_ = 4.4 x 10^-05^ s^-1^. This value was at least two orders of magnitude lower than the *k*_cat_ values for the human PTPs we tested, suggesting that the catalytic conversion of the substrate is slower for NMA1982 compared to many human PTPs. Interestingly, a comparably low activity has been reported for CDKN3, the human PTP that is most similar to NMA1982 [30].

**Fig 3.**
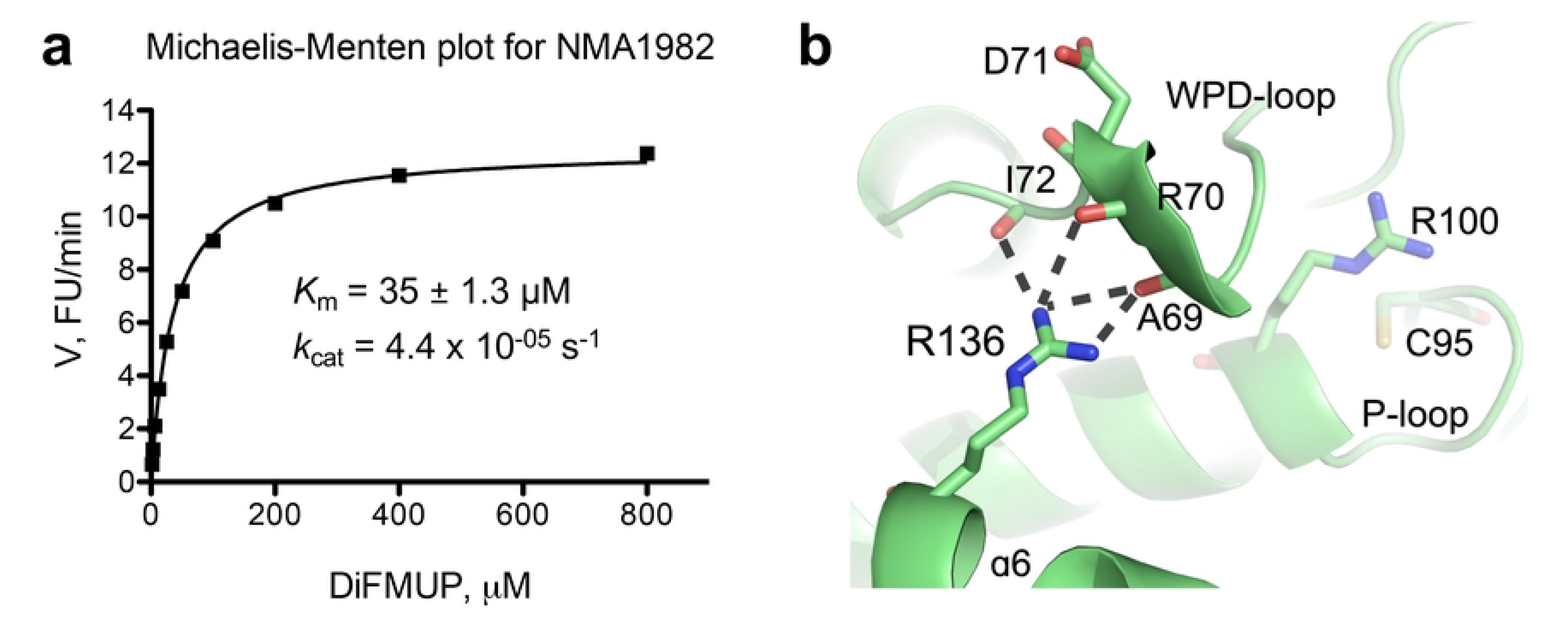
Assessing NMA1982 phosphatase activity. **(a)** Michaelis-Menten kinetics for NMA1982 using DiFMUP as the substrate. **(b)** Hydrogen bonding interactions between the side chain of R136 and the backbone oxygen atoms of WPD-loop residues A69, R70, and I72 in NMA1982 restrict WPD-loop dynamics.

### NMA1982 putative catalytic P-loop and WPD-loop mutagenesis studies

The putative P-loop in NMA1982, consisting of the **C95**-R-T-G-T-**R100** sequence, is one amino acid shorter than the typical PTP P-loop containing the **C**(X)_5_**R** signature motif. Superposition of the NMA1982 crystal structure with crystal structures of human DUSPs suggested that the putative catalytic cysteine C95 and the putative invariant arginine R100 in NMA1982 align very well with the corresponding residues in human DUSPs (shown for CDKN3 in **Fig 1b**). To test whether C95 and R100 are necessary for NMA1982 phosphatase activity, we generated and assessed the NMA1982 C95S and R100A mutant proteins. Since the corresponding residues are absolutely essential for enzymatic activity in all known PTPs [31], we expected the C95S and R100A mutants to be inactive. Indeed, both mutants showed no detectable phosphatase activity in the DiFMUP assay, even at protein concentrations as high as 60 µM (∼6-times higher than we used for assaying wt NMA1982). These data demonstrate that C95 and R100 of the C(X)_4_R motif are essential for NMA1982 phosphatase activity.

Next, we tested whether the putative catalytic acid/base D71 in the WPD-loop is critical for NMA1982 activity. Previous studies have shown that the mutation of the catalytic acid/base Asp to Ala typically results in a significant loss of phosphatase activity [32]. However, the degree of this loss in activity can vary widely among the PTPs and depends on whether other, nearby amino acids can compensate for the absence of Asp by providing a similar functionality [33]. To test the role of D71 for NMA1984 activity, we generated recombinant NMA1982 D71A mutant protein. We then assessed phosphatase activity of the mutant using a Michaelis-Menten kinetic assay with DiFMUP as the substrate. We found a substantial drop of 50% in the catalytic efficiency (*k*_cat_/*K*_m_) for the D71A mutant protein compared to wt NMA1982, demonstrating that D71 is important for catalysis (**Table 1**). Collectively, our data demonstrate that the putative catalytic amino acid residues in NMA1982 are indeed crucial for catalysis.

**Table 1.** Kinetic parameters of wild-type and mutant NMA1982 proteins.

| | K<sub>m</sub>, $\mu\text{M}$ | k<sub>cat</sub>, s<sup>-1</sup> | k<sub>cat</sub>/K<sub>m</sub>, M<sup>-1</sup>s<sup>-1</sup> |
| --- | --- | --- | --- |
| NMA1982 wt | 35 | 4.4×10<sup>-5</sup> | 1.26 |
| NMA1982 C95S | no activity detected |  |  |
| NMA1982 R100A | no activity detected |  |  |
| NMA1982 D71A | 40 | 2.7×10<sup>-5</sup> | 0.69 |
| NMA1982 R136A | 28 | $2.6 \times 10^{-4}$ | 9.28 |
| NMA1982 D71A/R136A | 33 | $1.1 \times 10^{-4}$ | 3.28 |

Given the relatively low activity we observed for wt NMA1982, we analyzed the NMA1982 crystal structure for additional insights Interestingly, we found that the side chain of R136 of the α6 helix, a residue that is unique to NMA1982 compared to human PTPs, forms multiple hydrogen bond interactions with the backbone oxygen atoms of A69, R70, and I72 of the WPD-loop (**Fig 3b**). Because restriction of the WPD-loop dynamics can result in decreased PTP activity [16-18], we hypothesized that the interactions between R136 and the WPD-loop residues could potentially interfere with WPD-loop dynamics, and thus limit enzymatic activity. To test this hypothesis, we generated the NMA1982 R136A mutant protein and measured its phosphatase activity using the DiFMUP assay. Notably, we found that the R136A mutant had a 7.4-times higher catalytic efficiency than wt NMA1982, and that this increase was mainly due to an increase in *k*_cat_ (**Table 1**). This suggested that the interactions of R136 with the WPD-loop residues limit turnover in NMA1982, most likely by restricting WPD-loop dynamics. To further support this hypothesis, we tested the effect of the D71A mutation on the activity of the R136A mutant compared to wt NMA1982. If the greater activity of R136A was due to an increase in WPD-loop flexibility, we expected that the D71A mutation would cause a greater drop in catalytic efficiency in the R136A mutant compared to wt. Indeed, the D71A mutation had a 6- times greater effect on catalytic turnover for the R136A mutant compared to wt NMA1982 (**Table 1**). Our results suggest that intramolecular interactions restrict WPD-loop mobility and therefore limit enzymatic activity of NMA1982. These data further demonstrate the importance of the WPD-loop for NMA1982 activity, and hence provide additional proof for NMA1982 utilizing a PTP catalytic mechanism.

### NMA1982 inhibition, pH and temperature dependence, and oxidative inactivation studies

To provide additional evidence that NMA1982 behaves like a typical PTP, we tested the enzyme’s response to sodium orthovanadate (Na_3_VO_4_), a general PTP inhibitor that resembles inorganic phosphate and binds into the catalytic pocket as a transition state analog [34]. Using our established DiFMUP assay, we found that orthovanadate inhibited NMA1982 activity in a dose-dependent manner with an IC_50_ value of 161 μM (**Fig 4a**). Thus, the potency of orthovanadate against NMA1982 was comparable to its potency against other human PTPs (Reference [35] and our unpublished data). Next, we tested the pH dependence of the NMA1982 phosphatase reaction. We subjected wt NMA1982 to catalytic rate measurements under various pH conditions and found that NMA1982 activity peaked at pH 6 (**Fig 4b**). These data agreed with a PTP peak activity that is typically observed between pH 5 and 6 [36]. Next, we tested how various temperatures affected NMA1982 activity. A close structural relative of NMA1982 is *Sso*PTP from *Sulfolobus solfataricus*, which exhibits peak phosphatase activity at 90°C [37]. As shown in **Fig 4c**, NMA1982 exhibited peak activity around 37°C, and activity was substantially reduced at 50°C, and completely absent at 80°C. Thus, NMA1982 was most active at temperatures at which eukaryotic PTPs show peak activity. Finally, we assessed NMA1982 activity in oxidative inactivation and rescue studies. Because PTP activity depends on the catalytic cysteine in the reduced state, and hydrogen peroxide treatment results in thiol oxidation and PTP inactivation [38], we tested whether NMA1982 activity was similarly sensitive to oxidation. Thus, we preincubated NMA1982 with various concentrations of hydrogen peroxide for 1 or 2 h and subsequently determined catalytic activity compared to a non-treated control (**Fig 4d**). Our data demonstrated that hydrogen peroxide treatment decreased NMA1982 activity, and that this decrease was time-and dose-dependent. Since the cysteine thiol can be oxidized to several oxidation states that vary in terms of reversibility with reducing agents such as dithiothreitol (DTT) [39], we also tested whether DTT treatment after peroxide treatment can rescue NMA1982 activity (**Fig 4e**). Our data showed that DTT treatment could partially rescue NMA1982 activity, indicating the presence of both thiol-reversible and -irreversible oxidation states. Higher peroxide concentrations (≥400 µM) and longer treatment tended to cause more irreversible oxidation of NMA1982. Collectively, our studies demonstrate that NMA1982 behaves like a typical PTP when subjected to a transition state inhibitor, various pH and temperature conditions, or oxidizing agents.

**Fig 4.**
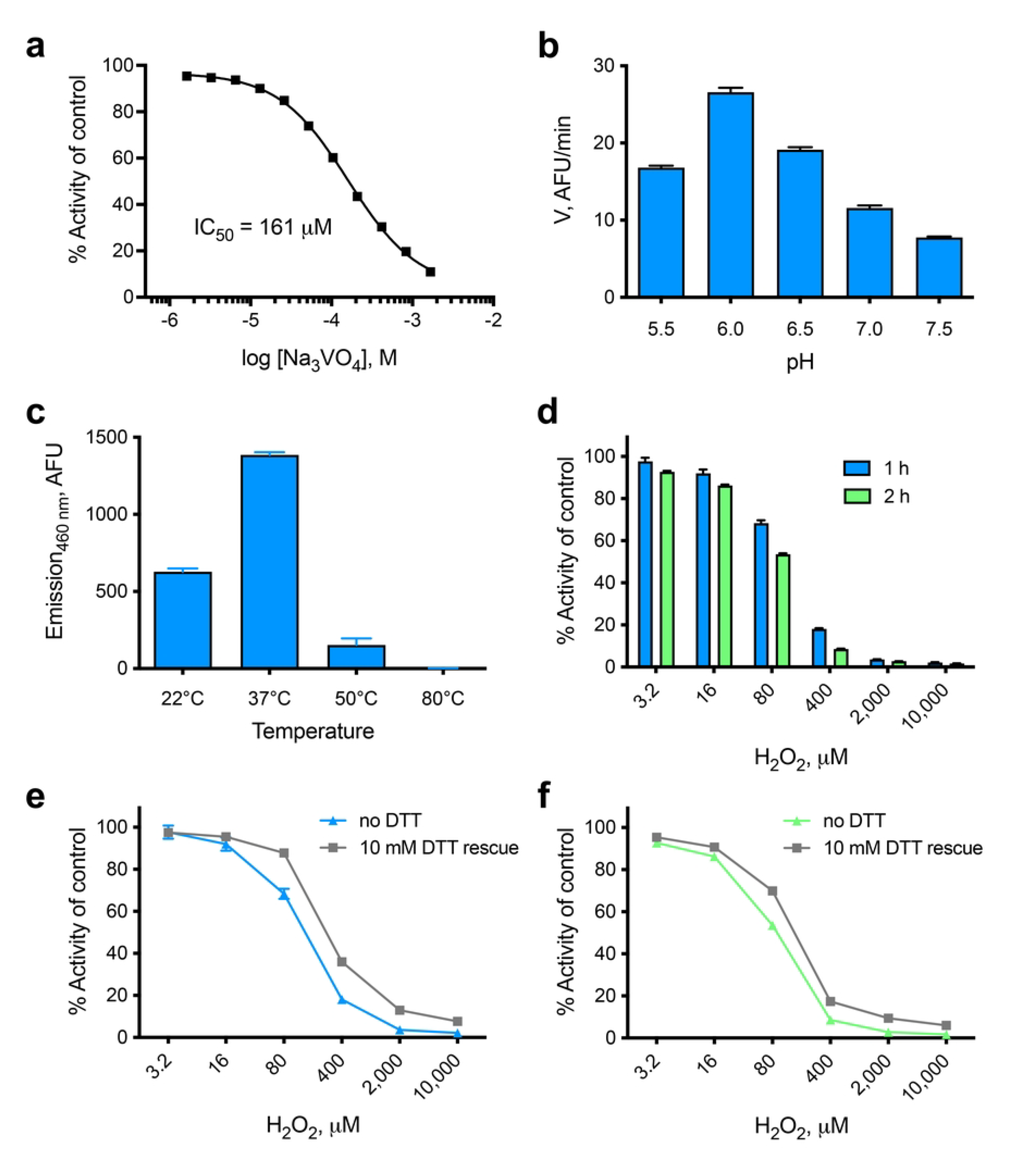
Biochemical characterization of NMA1982. **(a)** Dose-response inhibition of NMA1982 activity by the transition state-mimicking pan-PTP inhibitor sodium orthovanadate (Na_3_VO_4_). **(b)** NMA1982 activity, expressed as initial rate (V) in arbitrary fluorescence units (AFU) per minute, at various pH conditions as indicated. **(c)** NMA1982 activity at various temperatures as indicated. **(d)** Remaining NMA1982 activity in response to treatment with various concentrations of hydrogen peroxide for 1 or 2 h. **(e/f)** Remaining NMA1982 activity in response to treatment with various concentrations of hydrogen peroxide for 1 h (e) or 2 h (f) after rescue with 10 mM DTT or no DTT treatment. All data represent triplicate experiments and are presented as average ± SD.

### *N. meningitidis* NMA1982 secretion studies

PTPs encoded by bacteria have been shown to be secreted by many pathogens during infection, particularly by intracellular pathogens that can thereby directly target eukaryotic effectors [40]. Extracellular pathogens such as *N. meningitidis* are capable of secreting various proteins and virulence factors to promote their growth, for example, to allow biofilm formation or to target the host response [41, 42]. Furthermore, *N. meningitidis* can target host cells by secreting toxins that are endocytosed, such as the C2 fragment of the neisserial heparin-binding antigen (NHBA), which is released upon proteolysis [43], or PorB porin, which is released via outer membrane vesicles (OMVs) [44, 45]. The similarity of NMA1982 to eukaryotic PTPs strongly suggests that it may act as a virulence factor. We therefore assessed the secretion of NMA1982 in the model organism *N. meningitidis* 8013, which expresses the NMA1982 ortholog NMV0640 (**Fig S2**). A derivative of the wt NEM8013 strain lacking NMV0640 was engineered as a control (ΔNMV0640). Both wt and ΔNMV0640 strains were grown with agitation in rich medium before the cells and supernatant were separated, collected, and processed by SDS-Page and immunoblot analysis (**Fig 5**). The antibodies raised against NMA1982 clearly detected NMV0640 in both cell lysate and supernatant of the wt NEM8013 strain, whereas NMV0640 was not detected in the ΔNMV0640 control strain. As expected, PilE, the main component of type IV pili, was also recovered in the supernatant of the bacteria. The cytosolic marker NADP glutamate dehydrogenase and the outer membrane marker RMP4 were both absent from the supernatant fraction, confirming that there was no contamination of the supernatant fraction. Thus, our data demonstrate that *N. meningitidis* secretes NMA1982 during growth. Three secretion pathways are known to be active in *N. meningitidis*: the autotransporter, the two-partner secretion (TPS), and the type I secretion systems (T1S) [46]. The T1S is devoted to the secretion of the iron-regulated proteins FrpA/C only. Secretory proteins of the autotransporter and TPS first cross the inner membrane via the general secretion (Sec) or the twin arginine translocation (Tat) pathways. Periplasmic and outer membrane components also rely on the Sec and Tat systems to translocate through the cytosolic membrane. We therefore looked for a signal peptide in the sequence of NMA1982 that would indicate secretion by these pathways. Using both PrediSi [47] and SignalP6.0 [48], no known peptide signals were found, suggesting that NMA1982 is secreted via a Tat and Sec independent pathway.

**Fig 5.**
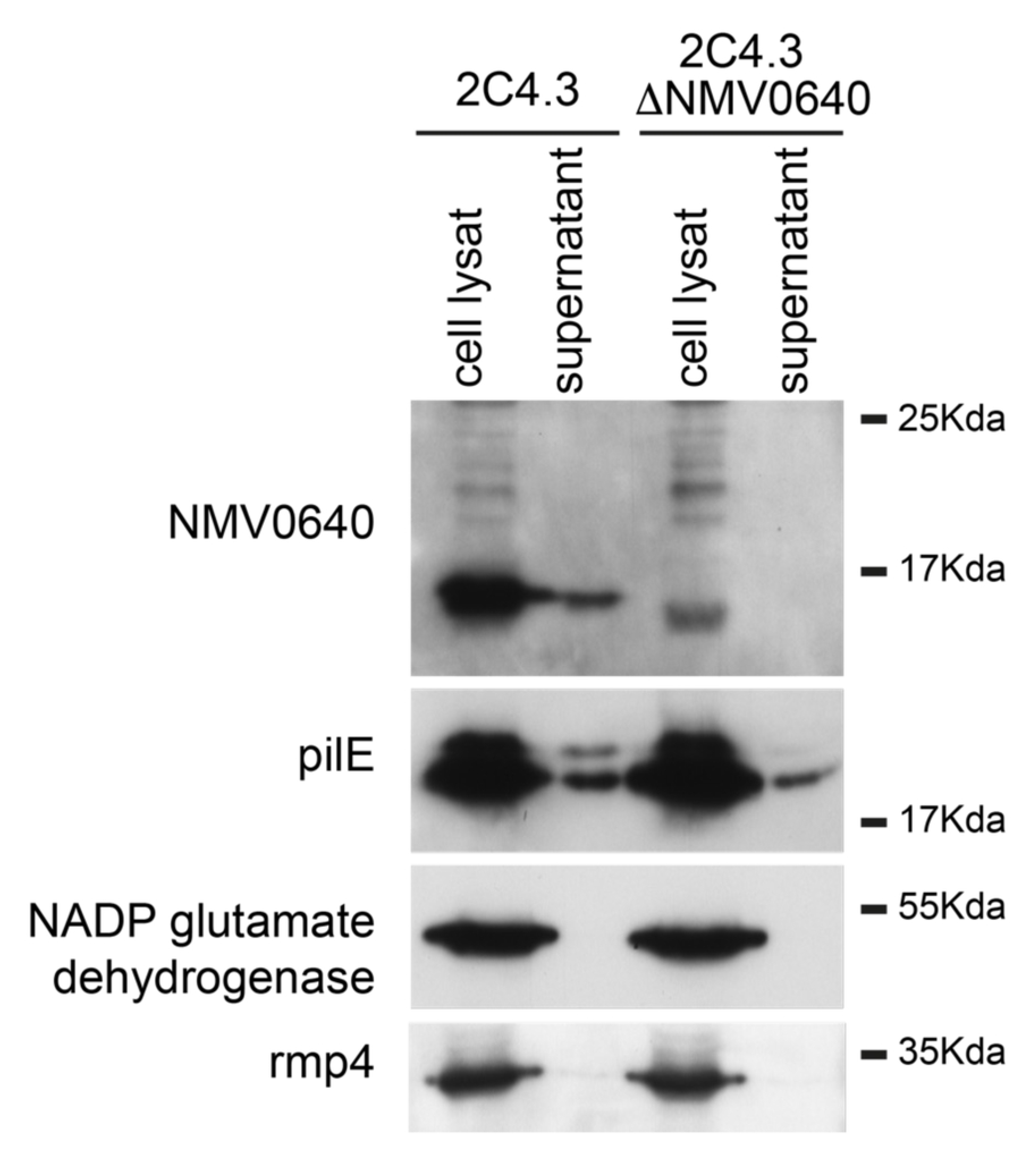
NMA1982 is recovered from the bacterial supernatant. Whole cell lysate and supernatant were analyzed by SDS-Page and immunoblotting. NMA1982 and PilE are present in the supernatant. PilE is used as a positive control since PilE is usually enriched in the bacterial supernatant. Rmp4 is an outer membrane protein thought to be enriched in outer membrane vesicles. Absence of Rmp4 in the supernatant suggests that the supernatants are not contaminated with membrane or vesicles. NADP glutamate dehydrogenase is a cytosolic enzyme used as a control. Interestingly, NADP glutamate dehydrogenase is recovered from a supernatant of an overnight culture of meningococci confirming that this protein is released during cell lysis (data not shown).

## Discussion

*N. meningitidis* causes systemic meningococcal disease, the mortality rate of which, even with optimal treatment, is still about 10% [21]. Here, we investigated an *N. meningitidis* protein that is highly similar to eukaryotic PTPs, enzymes that are considered crucial for many essential cellular processes. We show that NMA1982 uses a catalytic mechanism that is specific to PTPs, involving a highly conserved phosphate-binding loop that forms the catalytic center. This loop is usually seven amino acids in length and contains the hallmark C(X)_5_R signature motif. In NMA1982 the P-loop is shorter by one amino acid and contains a novel C(X)_4_R motif. Despite being shorter, the P-loop can accommodate a PTP substrate equally well as demonstrated by the similar *K*_m_ values of the pTyr mimetic DiFMUP for NMA1982 and human PTPs. In NMA1982, the catalytic cysteine and invariant arginine of the P-loop occupy space in the 3D structure that is similar to human PTPs. This ensures that the catalytic mechanism is not compromised. Given the existence of several bacterial NMA1982 homologs containing a C(X)_4_R motif (**Fig S3**), other phosphatases with that motif may exist in bacteria. Notably, a conserved glycine within the P-loop of eukaryotic PTPs is also 100% conserved in all bacterial homologs (G98 in NMA1982). The relatively low catalytic activity of NMA1982 is not unprecedented among the PTPs. The structurally closest human enzyme, CDKN3, exhibits a similarly low turnover number [30]. Interestingly, CDKN3 activity, which critically regulates cell cycle progression, is itself regulated by protein-protein interactions, resulting in a substantial increase of phosphatase activity under physiological conditions [49, 50]. Similarly, activity of NMA1982 might be enhanced *in vivo*. Our data demonstrates that NMA1982 is a PTP, enzymes that are essential for most cellular processes, not only in eukaryotes but also in bacteria. Importantly, we show that NMA1982, like known essential virulence factors, is secreted by *N. meningitidis*. The absence of a signal peptide suggests that NMA1982 is secreted by an atypical secretion machinery or via incorporation in OMVs as has been described for PorB. Future studies will need to address the question whether NMA1982 is indeed crucial for *N. meningitidis* survival and virulence, and thus may validate NMA1982 as a novel therapeutic target for the treatment of meningococcal diseases. Human PTPs have become attractive drug targets for many serious conditions, most notably cancer [51-53]. Compared to human PTPs, the shorter P-loop in NMA1982 results in a unique active site conformation, which may allow for the development of small molecule inhibitors with selectivity for the *N. meningitidis* protein.

## Acknowledgments

Research reported in this publication was supported by the National Institutes of Health under Awards Number 5U54-GM094586-02 (to L.T.) and NCI Cancer Center Support Grant P30CA030199. We thank Drs. Ian Wilson and Scott Lesley and the Joint Center for Structural Genomics (JCSG) for helpful discussions and for generously providing us with the NMA1982 plasmid. We thank Dr. Andrey Bobkov for his help with the DSC measurements. We also thank Dr. Douglas Sheffler for reading the manuscript.

## Supporting information captions

**Fig S1. Performance of the polyclonal rabbit anti NMA1982 antibodies (Abnova, Taiwan).** Antibody performance from two batches (Lot #11229 and Lot #11230) was tested against recombinant His-NMA1982 using both ELISA and immunoblot assays.

**Fig S2. Clustal Omega Alignment of NMA1982 orthologs in various *Neisseria meningitidis* and *Neisseria gonorrhoeae* strains.**

DNA sequences were obtained from https://mage.genoscope.cns.fr/microscope/home/index.php.

Start codons were confirmed using RBS Calculator (https://salislab.net/).

**Fig S3. Bacterial homologs of NMA1982.**

Clustal W alignment of NMA1982 and homologous protein sequences. Proteins other than NMA1982 are referred to by their NCBI Reference Sequence numbers. The ruler refers to NMA1982 residue numbers. The consensus (‘Majority’) and the consensus strength (‘+ Majority’) are given at the top. The color code of the consensus strength is red, 100% conserved, and dark blue, least conserved.

